# Assessment of gibberellin (GA_4+7_) mediated changes on grain filling, hormonal behaviour and antioxidants in high-density maize (Zea Mays L.)

**DOI:** 10.1101/511063

**Authors:** Wenwen Cui, Bingyun Zuo, Quanhao Song, Muhammad Kamran, Shahzad Ali, Qingfang Han, Zhikuan Jia

## Abstract

Dense plant cultivation is an efficient approach to improve the maize production by maximizing the utilization of energy and nutrient. However, dense plant populations may aggravate the abortion rate of young grains and result in fewer number of kernels per ear. Grain filling rate and duration play a decisive role in maize grain yield. Therefore, increasing plant density, consideration of enhancing the grain filling rate and duration of individual maize plant and regulating crop senescence would be the first priority. In this study, we examined the regulatory effects of GA_4+7_ under two application methods (shank-smearing and silk-smearing). Shank-soaking with GA_4+7_ at the rate of 0 (CK1), 10 (T1), 60 (T2), and 120 (T3) mg L^-1^, while silk-smearing at the rate of 0 (CK2), 10 (S1), 60 (S2), and 120 (S3) mg L^-1^ were used. The results showed that GA_4+7_ improved the grain filling rate by increasing the content of auxin, gibberellin and zeatin and abscisic acid in grains compared to control plants. In addition, The auxin, gibberellin and zeatin contents in the grains were positively and significantly correlated with the maximum grain weight and the maximum and mean grain-filling rates; the abscisic acid level was positively correlated with the maximum grain weight and the maximum and mean grain-filling rates. Moreover, GA_4+7_ increased the activities of superoxide dismutases, catalases, peroxidases, and reduced the malondialdehyde content in leaves compared with untreated plants. At the rate of 60 mg L^-1^, GA_4+7_ showed the greatest effect for shank-smearing and silk-smearing (T2 and S2), followed by 10 mg L^-1^ (T1) for shank-smearing and 120 mg L^-1^ (S3) for silk-smearing. Our results suggest that application of 60 mg L^-1^ GA_4+7_ for smearing application could efficiently be used for changed the level of hormones in grains and antioxidant enzymes in ear leaf, would be useful for enhancing grain filling rate and delaying leaves senescence, and resulting in an increasing of maize grain yield.

## 1 Introduction

Maize is one of the most important cereal crops and is widely used as food, fodder, and as industrial raw material worldwide [1]. Recently, the global production of maize has been exceeded that of rice and wheat [2]. Also, the global population is projected to further increase by 34% in 2050 (a total of 9.15 billion), thus an estimated 70% increase in agricultural production is demanded [3]. Achieving the food demands for such a huge population is a challenge of food security [4]. While, dense plant cultivation has the potential to attain higher crop productivity [5-8], which leads to greater leaf area index (LAI) and increases interception of photosynthetically available radiation (PAR), and enables the crops to use the intercepted solar radiation more efficiently [9-12]. However, dense plant cultivation should greatly affect the grain-filling process and result in lower maximum and average maize grain-filling rate [13-14]. Although, dense plant populations increase the number of spikes per unit area, but result in a decline of per-plant growth rates and exacerbate young kernel abortion, and so the ears per plant and kernels per ear were decreased [15-18]. Therefore, while adopting high planting density of maize, improving the individual maize grain filling rate is of great concern in modern crop systems.

Grain filling is the ultimate growth stage of cereals caryopses formation, which determines final weight of grains and thereby contribute greatly to grain productivity [19,20] The plant hormones have been stated to play a significant role in modifying grain filling progress and other various factors that regulated the grain filling progress. Yang, et al. [21] have reported that the wheat grain-filling rate is mediated by the balance between abscisic acid and ethylene, and the grain-filling rate increases with an increase in the ratio of ABA to ethylene in grains. The zeatin (Z) and zeatin riboside (ZR) in developing seeds have shown to temporarily improve endosperm cell division and fertilization during kernel setting [22]. The contents of indole-3-acetic acid (IAA) and abscisic acid (ABA) were higher in superior grains than in inferior [23], and the increased ABA and reduced IAA shortened grain-filling period in the inferior grains [24]. Liu, et al. [25] and Ali, et al. [26] have reported a positive and significant correlation of ABA, IAA, and Z + ZR contents with the maximum grain weight and grain-filling rates. In addition, a negative and significant correlation of ETH concentration with grain weight and grain-filling rates was reported by Liu, et al. [25]. Additionally, the higher gibberellins (GAs) contents were present at early stages of grain filling [27]. These studies showed that cereals grain filling is markedly affected by alterations in hormones level in grains.

In recent decades, plant growth regulators (PGRs) including gibberellins (GAs) are gaining interest among agriculture scientists are broadly used in agronomic crops. GAs comprise a large family of hormones that are ubiquitous in higher plants and have long been known as endogenous plant growth regulators, promoting several aspects of plant growth and developmental processes, such as cell division, stem elongation, seed germination, dormancy, leaf expansion, flower and fruit development [28,29]. They were first discovered in 1930s by Teijiro Yabuta and Yusuke Sumiki and named after the pathogenic fungus called *Gibberella fujikuroi* culture [30]. Since then 136 different types of gibberellins structures have been identified and isolated from different fungi and plants sources [31,32]. Despite a large number of GAs, relatively a few gibberellins such as GA1, GA3, GA4, GA7 are believed to have intrinsic biological activity in higher plants [29] (chemical structure of GAs with high bioactivity shown in Scheme 1). GA3 is one of the most widely used plant growth hormones. With the deepening of research, other hormones of GAs, especially GA4 and GA7 attract more and more attention due to their special effects on plants. Some researchers suggested that GA_4+7_ could successfully induce the fruit set and increased fruit size in cucumber [33], pear [34], apple [35]. In addition, compared to pollinated fruit, GA_4+7_-treated fruits accumulated more quantities of sucrose and fewer organic acids [34]. Therefore, the application of GA_4+7_ proved to be an utmost agricultural remedy in horticultural crops, which would otherwise dramatically increase yield and income. However, the application of GA_4+7_ in cereal crops is limited and the effect of exogenous GA_4+7_ on the regulation of maize grain filling and its mechanism remains unclear.

**Scheme 1.**
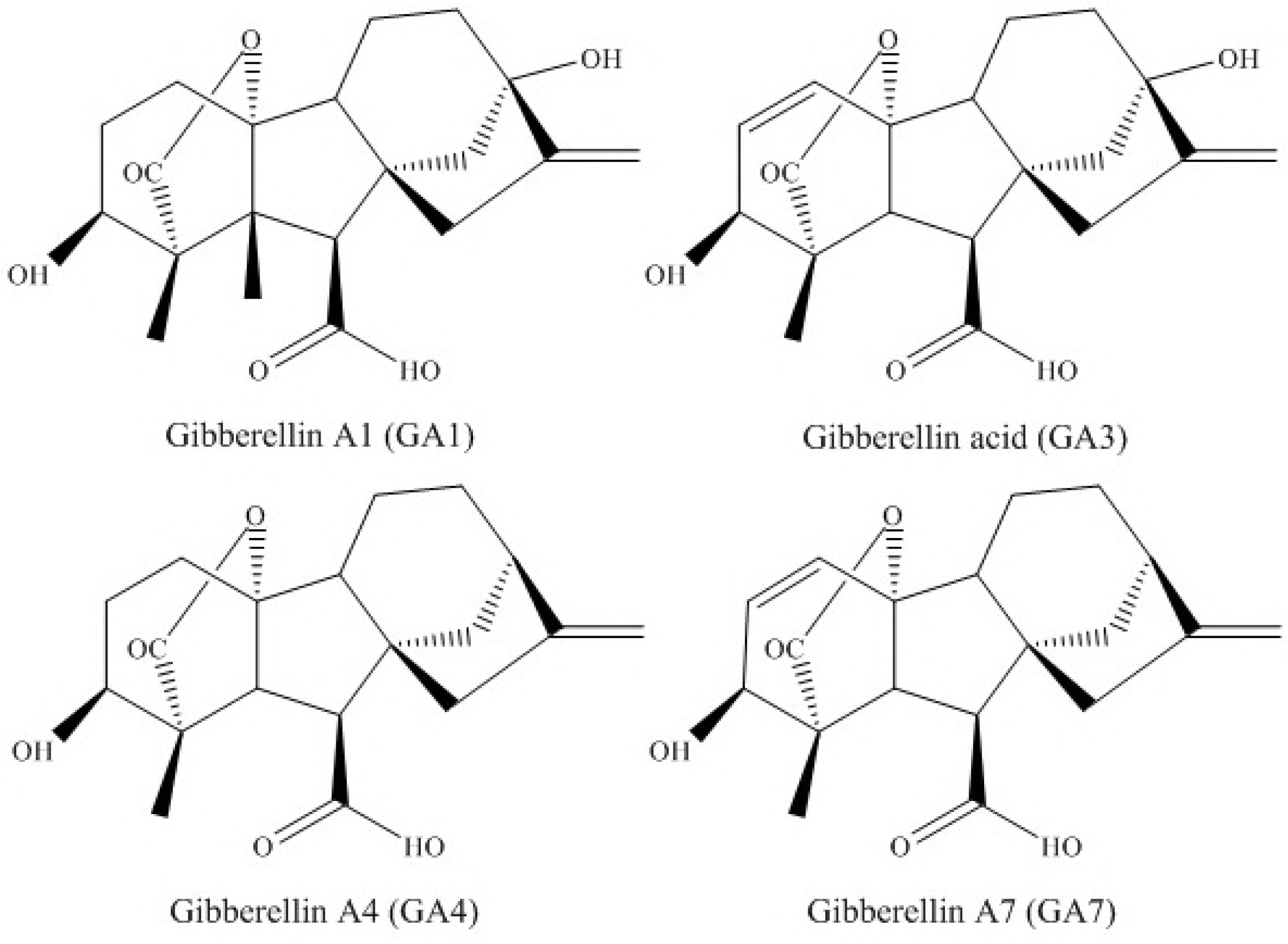
The chemical structure of GAs with high bioactivity

Reactive oxygen species (ROS) are continually being produced in plants, which can act as a signal molecule in plants and trigger a series of cellular responses [36,37]. In plants, an increase in the ROS levels exceeding the detoxification levels of plant tissues could be toxic. However, plants have evolved an enzymatic and non-enzymatic antioxidants defense mechanism to effectively scavenge the ROS and maintain a proper balance within the plants. Enzymatic antioxidants mainly include superoxide dismutase (SOD) peroxidase (POD) ascorbate peroxidase (APX), and catalase (CAT). However, senescence and various environmental stresses could disrupt the balance between ROS generation and detoxification, resulting in lipid peroxidation, chlorophyll degradation and loss of cell membrane integrity [38]. The research of exogenous GA_4+7_ stress on antioxidants in maize is limited and unclear.

These previous literature have indicate that the potential of GA_4+7_ to control grain filling of maize is limited, and the possible effects on hormone change and antioxidants have not been investigated in details. Therefore, the objectives of the present study were to instigate the effects of shank-smearing and silk-smearing with different concentrations of GA_4+7_ on grain filling rate, hormonal changes and its relation with grain filling and possible changes in antioxidants. We provided the first detailed description of the GA_4+7_ application in improving grain filling rate of maize and antioxidants, with the aim of achieving higher grain yield in high-density maize.

## 2 Materials and methods

### 2.1 Seed and reagent source

The hybrid maize seeds (Zhengdan, 958, a local hybrid) were provided by China National Seeds Group Co., LTD. The seed moisture content and germination potential were 13% and 90% respectively. The test reagent was Gibberellin GA_4+7_ [*m* (GA4): *m* (GA7) = 40:60] (BR, purity ≥ 90.0%) was purchased from Shanghai Ryon Biological Technology Co., Ltd, China.

### 2.2 Study site description

Field trials were conducted in 2015-2016 at the Institute of Water Saving Agriculture in Semi-Arid Regions of China, Northwest A&F University, Yangling (34°20′N, 108°04′E, 454.8 m altitude), Shaanxi Province, China. The climate is temperate semi-arid monsoon with a mean annual temperature of 12-14 °C, and the means annual precipitation and evaporation were 580.5 mm and 993.2 mm, respectively. The total yearly sunshine duration was 2158 h, and the no-frost period was 221 days. Most of the rainfall occurred in hot seasons from July to September (Fig.1). The soil of the experimental site consists of Cumuli-Ustic Isohumosols (Chinese Soil Taxonomy), and the soil characteristics from 0 to 40 cm deep at the research site is shown in Table 1.

**Fig. 1.**
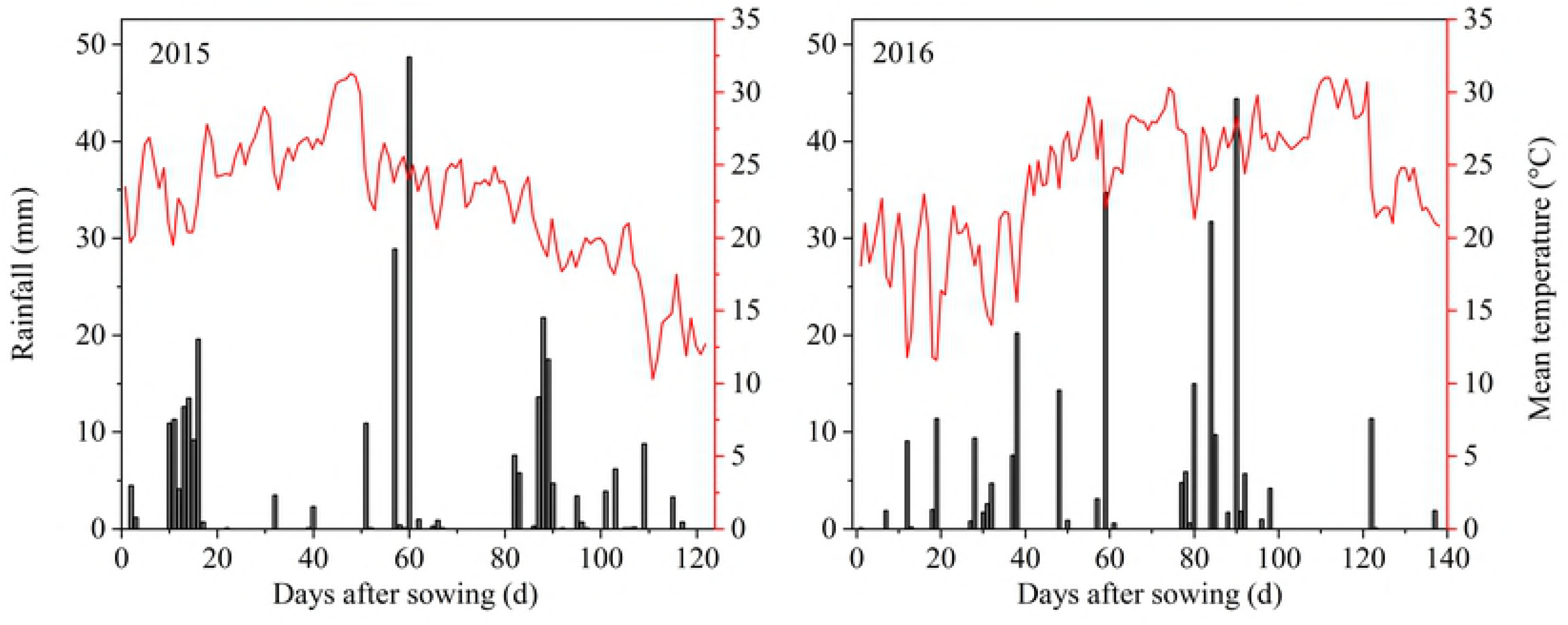
Mean temperature and rainfall during the growing seasons in 2015 and 2016.

**Table 1.**
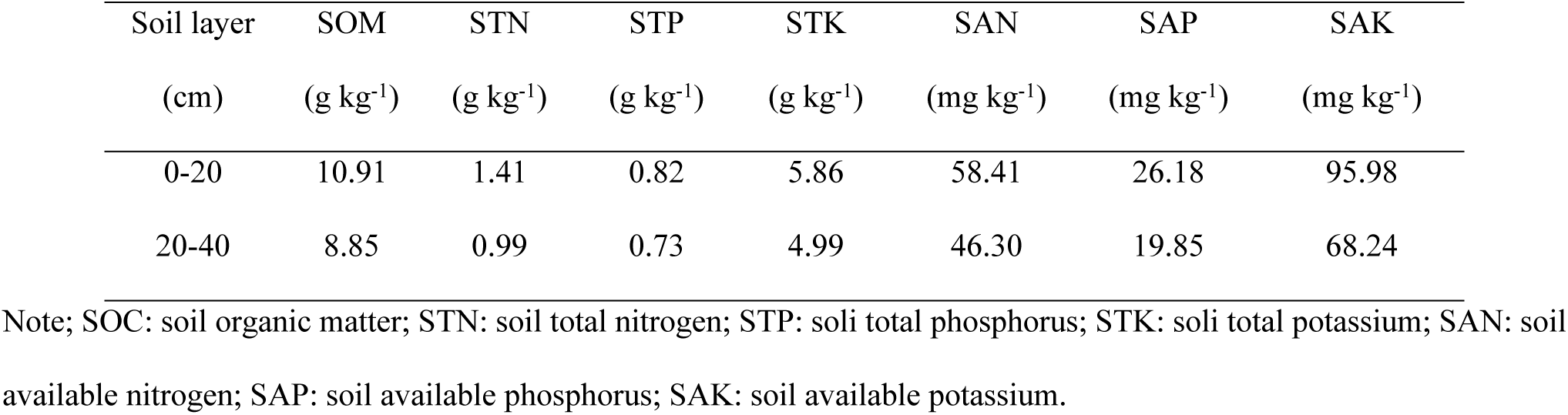
Soil chemical properties of the top 40 cm in the experimental fields at the Institute of Water Saving Agriculture in Semi-Arid Regions of China in Northwest A&F University, Yangling, Shannxi Province, China, in 2015 and 2016.

| Soil layer | SOM | STN | STP | STK | SAN | SAP | SAK |
| --- | --- | --- | --- | --- | --- | --- | --- |
| (cm) | (g kg<sup>-1</sup>) | (g kg<sup>-1</sup>) | (g kg<sup>-1</sup>) | (g kg<sup>-1</sup>) | (mg kg<sup>-1</sup>) | (mg kg<sup>-1</sup>) | (mg kg<sup>-1</sup>) |
| 0-20 | 10.91 | 1.41 | 0.82 | 5.86 | 58.41 | 26.18 | 95.98 |
| 20-40 | 8.85 | 0.99 | 0.73 | 4.99 | 46.30 | 19.85 | 68.24 |

Note; SOC: soil organic matter; STN: soil total nitrogen; STP: soli total phosphorus; STK: soli total potassium; SAN: soil available nitrogen; SAP: soil available phosphorus; SAK: soil available potassium.

### 2.3 Experimental design and treatments

The regulatory effect of GA_4+7_ under two application methods (shank-smearing and silk-smearing) were studied in 2015-2016. Shank-smearing with GA_4+7_ at the rate of 0 (CK1), 10 (T1), 60 (T2), and 120 (T3) mg L-1, while silk-smearing also at the rate of 0 (CK2), 10 (S1), 60 (S2), and 120 (S3) mg L-1 were used. Smearing applications were conducted at one week after silking stage (R1). The experiment was laid out in a randomized complete block design (RCBD) using three replications. Maize seeds were manually sown at a density of 97,500 plants ha^-1^ on 13 and 25 June 2015 and 2016, and harvested on 14 and 21 October 2015 and 2016, respectively. The plot size was 45 m^2^ (5m x 9m) with the plant-plant and row-row distance of 17 and 60 cm, respectively. All plots were supplied with 225 kg N ha^-1^, and 120 kg P_2_O_5_ ha^-1^. All P fertilizers and 60% of the N fertilizer were applied at pre-sowing. The remaining 40% of urea (N 46%) was applied as a top dressing at the twelfth leaf stage (V12). During the entire growth period, disease, pest and weed control in each treatment were well controlled. Irrigation was applied when it was necessary.

### 2.4 Sampling and measurements

At tasseling stage (VT), healthy and uniform maize plants were marked in each plot. Three marked ear leaves from each plot were sampled at 10 days intervals from the 15 days after silking (DAS) to maturity. Ear leaves were stored in liquid nitrogen for the determination of reactive oxygen species. While three marked ears from each plot were sampled at 7 days intervals from the 14 DAS to maturity. The averages grain weight was taken for the three representative ear samples from each time point. First, 100 grain was sampled from the middle part of the maize ear. Half of the sampled grains were used for the hormones measurements, and remaining grains were oven dried at 70 °C until to a constant weight.

#### 2.4.1 Grain-filling process

The grain-filling data were fitted using the Richards growth equation [39]:

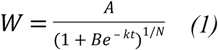

The grain-filling rate (G) was calculated as the derivative of *Eq* (1):

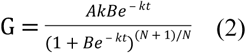

Where W is the grain weight (mg), A is the final grain weight (mg), t is the time after anthesis (d), and B, k and N are coefficients determined by regression analysis.

#### 2.4.2 Endogenous hormone

The extraction, purification, and quantification of IAA, ZR, GA_3_, ABA by an indirect ELISA technique were essentially identical to those described by Liu, et al. [25] and Yang, et al. [27]. The antigens and monoclonal antibodies against IAA, ZR, GA_3_, ABA and immunoglobulin G-horseradish peroxidase (IgG-HRP) used in the ELISA were produced at the Phytohormones Research Institute, China Agricultural University, China.

For the extraction and purification of IAA, ZR, GA_3_ and ABA, a sample of approximately 0.5 g was ground in ice bath with 4 mL 80% (v/v) methanol extraction buffer containing 1 mmol L^-1^ butylated hydroxytoluene (BHT) as an antioxidant. The homogenates were incubated at 4 °C for 4 h and centrifuged at the same temperature. Afterwards the supernatants were taken into a new centrifuge tube, precipitated with 1 mL methanol extraction buffer, incubated for 1 h and centrifuged again. The supernatants were passed through Chromosep C18 columns (C18 Sep-Park Cartridge, Waters Corp, USA) and prewashed with 5 mL 100% ether and then 5 mL 100% methanol. The hormone fractions were dried with N_2_ and dissolved in 1 mL phosphate-buffered saline (PBS) containing 0.1% (v/v) Tween-20 and 0.1% (w/v) gelatin (pH 7.5) for enzyme-linked immunosorbent assay (ELISA) evaluation.

The quantification of IAA, ZR, GA_3_ and ABA was done using 50 µL sample (standard sample or the test sample) and 50 µL of the diluted antibody were added into each well of the elisa plate, and the plate was placed in the wet box at 37 °C for 0.5 h. The plate was then washed 4 times with phosphate-buffered saline (PBS) containing 0.1%(v/v) Tween-20, and Then add 100 µL of the diluted IgG-HRP was added to each well, and the plate was put in the wet box at 37 °C for 0.5 h. Afterwards, the plate was washed 4 times again, and then it's the color reaction. To each well will be added 100 µL coloration liquid containing 0.1% (W/V) o-phenylenediamine (OPD) and 0.04% (V/V) H_2_O_2_ in substrate buffer (the substrate buffer contains 5.10g C_6_H_8_O_7_·H2O, 18.43g Na_2_HPO_4_·12H_2_O and 1 mL Tween-20 per litre with pH 5). The chromogenic reaction was also carried out in the wet box, and the reaction was terminated by adding 50 µL 2 mol L^-1^ H_2_SO_4_. The absorbance of the solution was read at 490 nm by a spectrophotometer.

The logit curve was used to calculate the ELISA results. Hormone concentration was expressed as ng g^-1^ fresh weight.

#### 2.4.3 Antioxidant enzyme

For enzyme extraction, 0.5 g of fresh leaf samples were ground with 5mL pre-cooled 50 mM phosphate buffer (pH 7.5). The homogenates were centrifuged at 13000 rpm for 30 min at 4 °C, and the resulting supernatants were used for enzyme assay. All enzyme activity data were expressed as U mg^-1^ fresh weight (FW).

Superoxide dismutase (SOD) activity was determined by recording the decrease in the absorbance of nitro-blue tetrazolium (NBT) by the enzyme [40]. The reaction mixture consisting of 1.5 mL 50 mM phosphate buffer (pH 7.8), 0.3 mL 130 mmol L^-1^ methionine, 0.3 mL 750 mol L^-1^ NBT, 0.3 mL 100 mol L^-1^ EDTA-Na_2_, 0.3 mL 20 mol L^-1^ FD and 0.3 mL distilled water was mixed with 20 μL crude enzyme extract. Control and enzyme solution was placed at 4000 lux light for 30 min. The blank sample was placed in the dark. The absorbance of the solution was monitored at 560 nm by an ultraviolet spectrophotometer.

Peroxidase (POD) activity was assayed according to Ekmekci and Terzioglu [41]. 20 μL enzyme extract was drawn and mixed with 3mL of POD reaction solution. The reaction mixture contained 1.5 mL 50 mM sodium phosphate buffer (pH 6.0), 0.5 mL H_2_O_2_ (30%), 0.5 mL 50 mMguaiacol, and 0.5 mL distilled water. The absorbance values were recorded once every 30 s at 470 nm by using an ultraviolet spectrophotometer.

Catalase (CAT) activity was assayed by measuring the decomposition of H_2_O_2_ [42]. The reaction mixture contained 1.5 mL 50 mM sodium phosphate buffer (pH 7.0), 0.2 mL crude extract, 1 mL distilled water and 0.3 mL 100 mM H_2_O_2_. The breakdown of H_2_O_2_ was followed by measuring the absorbance change at 240 nm.

The lipid peroxidation was determined by measuring the content of malondialdehyde (MDA) by following the procedure of Zhang [43]. 0.5 g leaf samples were homogenized in 5 mL of 5 % trichloroacetic acid (TCA), and the homogenate was centrifuged at 4000 g for 10 min at 25 °C. 2 mL enzyme solution was drawn and mixed with 3 mL of 2-thiobarbituric acid (TBA) in 20 % trichloroacetic acid. The mixture was water-bath heated at 100°C for 20 min and centrifuged for cooling down. The absorbance of the supernatant was determined respectively at 450 nm, 532 nm, and 600 nm.

#### 2.4.4 Yield and yield components

To determine ear characteristics and grain yield, thirty representative plants from each replicate were sampled at physiological maturity. Kernel number ear^-1^ and thousand kernels weight (TKW) were measured on 20 representative ears per plot. TKW was determined after drying a thousand grains at 75 °C to constant weight. Grain yield was determined at 13.0% moisture content.

### 2.5 Statistical analysis

Analyses of variance (ANOVA) were determined by using SAS 9.2 software (SAS Institute, Cary, NC, USA). The data from each sampling were analyzed separately. Least significance difference test (LSD) was applied to determine the significance between different treatments (p < 0.05).

## 3 Results

### 3.1 Effects of GA_4+7_ on yield and yield component

The results of field experiments in 2015-2016 showed that GA_4+7_ treatments significantly (P<0.05) enhanced the ear characteristics (ear length and diameter, kernels ear^-1^, and 1000 kernel weight) and grain yield of maize when compared with control treatments (Table 2). Compared with control treatment, T2 and S2 treatments had the best effects on the maize yield, followed by T1 and S3. The shank smearing treatments affected the grain yield mainly by increasing the thousand grain weight of maize, two years average thousand grain weight under T1 and T2 treatments were increased by 24.5g, 36.99 g, respectively, i.e. 8.5%, 12.8%. While the silk smearing treatments mainly increased kernel number to improve grain yield, two-year average kernel number under S2 and S3 treatments were increased by 77, 44 kernels per ear, respectively, i.e. 26.9% and 15.5%. In 2015, the highest concentration of GA_4+7_ in shank smearing treatment and the lowest concentration of GA_4+7_ in silk smearing treatment, i.e., T3 and S1 had a relatively small impact on increasing yield compared with control treatment (CK). However, the grain yield of T3 and S1 were still improved by a relatively large amount in 2016. The two year average grain yields under T1, T2, and T3 treatments were increased by 1050.4kg ha^-1^, 2132.6 kg ha^-1^, 348.8 kg ha^-1^, 12.1%, 24.6%, and 4.0% increases compared with CK1, whereas that of S1, S2, and S3 treatment were greater by 431.9 kg ha^-1^, 2863.4 kg ha^-1^, 1745.6 kg ha^-1^, respectively, i.e., 5.11%, 33.9%, and 20.7% increases compared with CK2.

**Table 2.** Effects of GA_4+7_ on ear length, ear diameter, kernels ear^-1^, thousand grain weight, and grain yield (t ha^-1^) of maize in 2015-2016.

|  | Treatments | Kernel number | Ear Length | Ear Diameter | Thousand kernel weight | Grain Yield |
| --- | --- | --- | --- | --- | --- | --- |
|  |  | (No.ear<sup>-1</sup>) | (cm) | (mm) | (g) | (t ha<sup>-1</sup>) |
| 2015 | CK1 | 477±7ab | 14.1±0.4b | 45.6±0.4b | 284.1±7.1b | 8.3±0.3c |
|  | T1 | 474±18ab | 14.9±0.2a | 46.2±1.1b | 311.9±9.6a | 9.5±0.3b |
|  | T2 | 501±12a | 14.9±0.1a | 49.7±0.7a | 319.1±4.1a | 10.5±0.7a |
|  | T3 | 470±17b | 14±0.5b | 45.3±0.9b | 294.2±8.8b | 8.9±0.5bc |
|  | CK2 | 475.1±2.3b | 14.1±0.5b | 45.8±0.1b | 285.1±6.2c | 8.2±0.22c |
|  | S1 | 462.1±25.2b | 14.1±0.3b | 45.3±0.7b | 297±2b | 8.16±0.49c |
|  | S2 | 523±20.1a | 15.4±0.5a | 47.1±0.5a | 311.7±6.5a | 10.88±0.14a |
|  | S3 | 489.2±19.9ab | 14.6±0.3ab | 46.8±0.3a | 305.4±6.2ab | 9.89±0.17b |
| 2016 | CK1 | 512±2c | 15.4±0.2c | 46.8±1b | 294.1±4.6c | 8.8±0.5c |
|  | T1 | 582±4b | 16.3±0.2b | 49.5±0.5a | 315.3±3.9b | 9.9±0.5b |
|  | T2 | 602±10a | 17.3±0.7a | 50.2±0.4a | 329.8±6.5a | 11.1±0.1a |
|  | T3 | 577±3b | 16.7±0.5ab | 48±0.9b | 312±3.9b | 9.1±0.4bc |
|  | CK2 | 506±6c | 15.2±0.5b | 46.9±0.5c | 290.8±7.5b | 8.7±0.03d |
|  | S1 | 581±6b | 17.2±0.1a | 49.7±0.2a | 310.3±7.1a | 9.6±0.02c |
|  | S2 | 613±9a | 17.5±0.4a | 49.3±0.6a | 320.9±3.9a | 11.74±0.76a |
|  | S3 | 581±7b | 17.4±0.2a | 48.3±0.5b | 307.7±9.6a | 10.5±0.05b |

CK1, T1, T2, and T3 represent shank-smearing with GA<sub>4+7</sub> at the rate of 0, 10, 60, and 120 mg L<sup>-1</sup>, respectively. CK2, S1, S2 and S3 represent silk-smearing with GA<sub>4+7</sub> at the rate of 0, 10, 60, 120 mg L<sup>-1</sup>, respectively. Data are expressed as the mean ± S.D. (n=3). Means followed by different letters within a column are significantly different at P <0.05 as determined by the LSD test.

### 3.2 Effects of GA_4+7_ on grain filling rate

GA_4+7_ treatments showed positive effects on grain filling of maize (Fig.2 and Table 3). The grain filling rate reached was maximized at 28 DAS for all treatments. The maximum grain filling rate under T1, T2, and T3 treatments were increased by 0.674mg grain^-1^ d^-1^, 2.399 mg grain^-1^ d^-1^, 0.609 mg grain^-1^ d^-1^, 7.9%, 28.3%, and 7.2% increases compared with CK1, whereas that of S1, S2 and S3 treatments were greater by 0.429mg grain^-1^ d^-1^, 1.194 mg grain^-1^ d^-1^, 0.633 mg grain^-1^ d^-1^, respectively, i.e., 5.02%, 14.0%, and 7.4% increases compared with CK2. GA_4+7_ application at the rate of 60 mg L^-1^ (S2 and T2 treatments) evidently improved the grain weights and the maximum and mean grain filling rates, compared to the control and other treatments. Compared with control, the grain filling rate in T2 treatment was increased by 32.7%, 27.8%, 28.3%, 11.5%, 11.1% and 49.0% at 14, 21, 28, 35, 42, 49 DAS, while S2 treatment was increased by 18.2%, 22.8%, 14.0%, 10.9%, 15.8% and 37.4% at the respective growth stages. The grain filling rate and grain weight of T1 and S3 at all stages were only inferior to that of T2 and S2. Conversely, T3 and S1 had no significant effects on the maximum grain filling rate but had a significant effect on the mean grain filling and the maximum grain weight compared with control treatments.

**Fig. 2.**
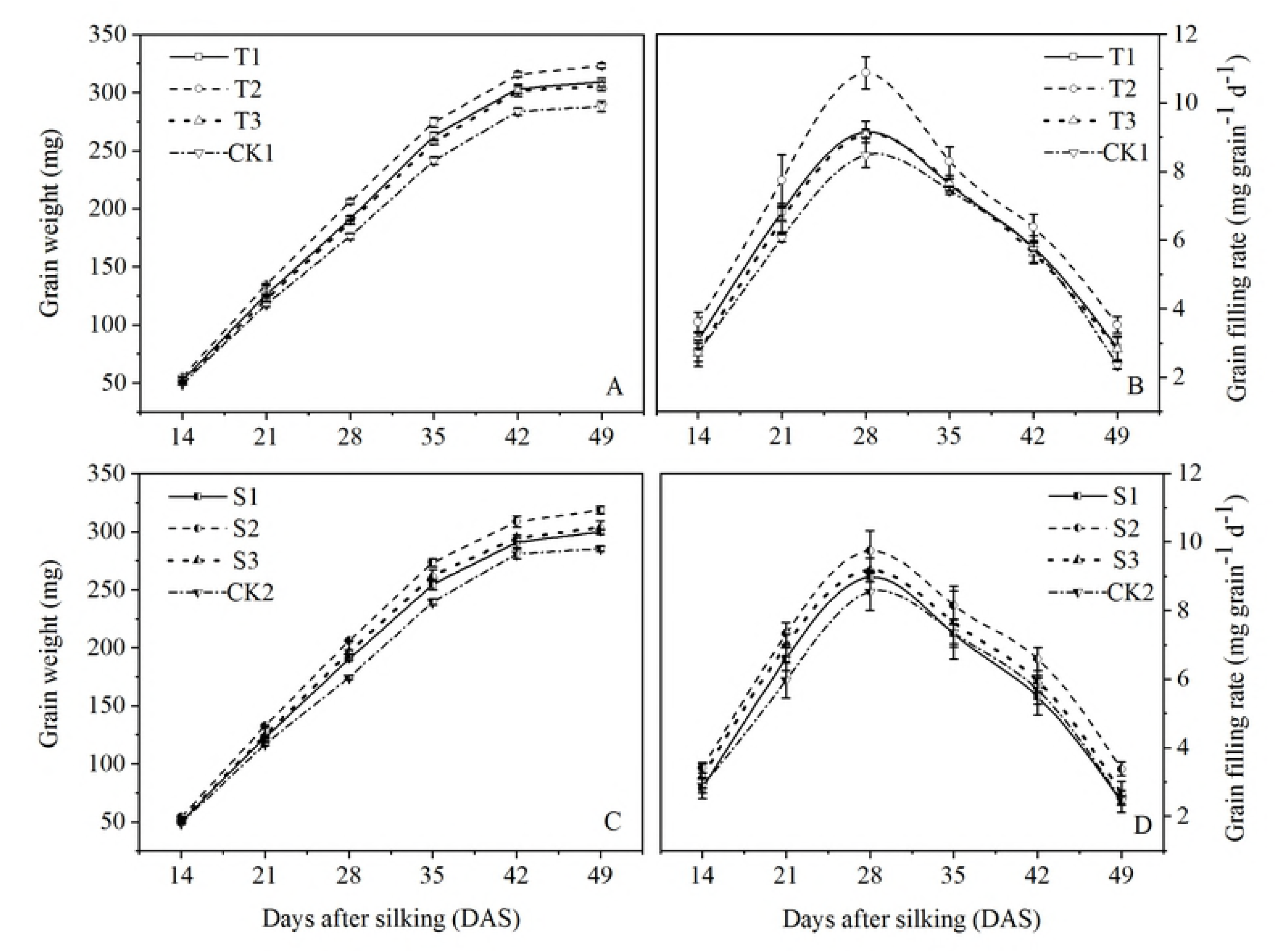
The effects of GA_4+7_ smearing application on grain weights and grain-filling rates of maize. CK1, T1, T2 and T3 represent shank-smearing with GA_4+7_ at the rate of 0, 10, 60, and 120 mg L^-1^, respectively. CK2, S1, S2 and S3 represent silk-smearing with GA_4+7_ at the rate of 0, 10, 60, 120 mg L^-1^, respectively. The vertical bars represent the ± the standard error of the mean (n = 3).

**Table 3.** Grain-filling characteristics of maize under different GA_4+7_ smearing application

| Treatments | W<sub>max</sub> (mg) | G<sub>max</sub> (mg grain<sup>-1</sup> d<sup>-1</sup>) | G<sub>mean</sub> (mg grain<sup>-1</sup> d<sup>-1</sup>) |
| --- | --- | --- | --- |
| CK1 | 288.3±4.51c | 8.48±0.37c | 5.47±0.08c |
| T1 | 309.0±3.81b | 9.16±0.31b | 5.89±0.15b |
| T2 | 323.2±2.2a | 10.88±0.47a | 6.74±0.05a |
| T3 | 305.8±3.86b | 9.09±0.12bc | 5.75±0.11b |
| CK2 | 285.3±2.18c | 8.55±0.54b | 5.48±0.24c |
| S1 | 300.1±2.09b | 8.98±0.14ab | 5.61±0.03bc |
| S2 | 318.5±3.09a | 9.74±0.58a | 6.43±0.06a |
| S3 | 303.6±5.64b | 9.18±0.35ab | 5.93±0.37b |

CK1, T1, T2 and T3 represent shank-smearing with GA<sub>4+7</sub> at the rate of 0, 10, 60, and 120 mg L<sup>-1</sup>, respectively. CK2, S1, S2 and S3 represent silk-smearing with GA<sub>4+7</sub> at the rate of 0, 10, 60, 120 mg L<sup>-1</sup>, respectively. Data are expressed as the mean ± S.D. (n=3). Means followed by different letters within a column are significantly different at P < 0.05 as determined by the LSD test. W<sub>max</sub>: the final grain weight; G<sub>max</sub>: maximum grain-filling rates; G<sub>mean</sub>: mean grain-filling rates.

### 3.3 Effect of GA_4+7_ on hormonal changes in grains

#### 3.3.1 IAA and ZR contents in grains

The IAA and ZR contents in maize grains exhibited a similar pattern during the grain filling process. The IAA and ZR contents increased linearly during the initial grain-filling stage and attained the maximum peak curves at 28 DAS for all treatments (Fig. 3 and 4). The IAA and ZR contents under shank-smearing treatments were greater than that under CK1 from 14 to 28 DAS. Compared with CK1, the average IAA contents from 14 to 28 DAS under T1, T2 and T3 were increased by 50.2%, 82.7%, and 50.7%, while the average ZR contents from 14 to 28 DAS increased by 23.6%, 41.0%, and 14.6% under T1, T2 and T3 treatment. The IAA contents in grains under S2 and S3 treatments were greater than that under CK2 from 14 to 49 DAS. Compared with CK2, the mean IAA contents at all sampling stages of S2 and S3 were raised by 32.4% and 20.3%. At 49 DAS, S1 treatments have no significant effects on ZR contents in maize grains; however, the ZR contents in the grains under other shank-smearing and silk-smearing treatments in every sampling period were significantly higher than that under CK1 and CK2. And the mean ZR contents under T1, T2, and T3 treatments were increased by 25.5%, 45.2%, 24.0% compared with CK1, while that of S1, S2 and S3 increased by 20.7%, 37.0%, and 22.0% compared with CK2.

**Fig. 3.**
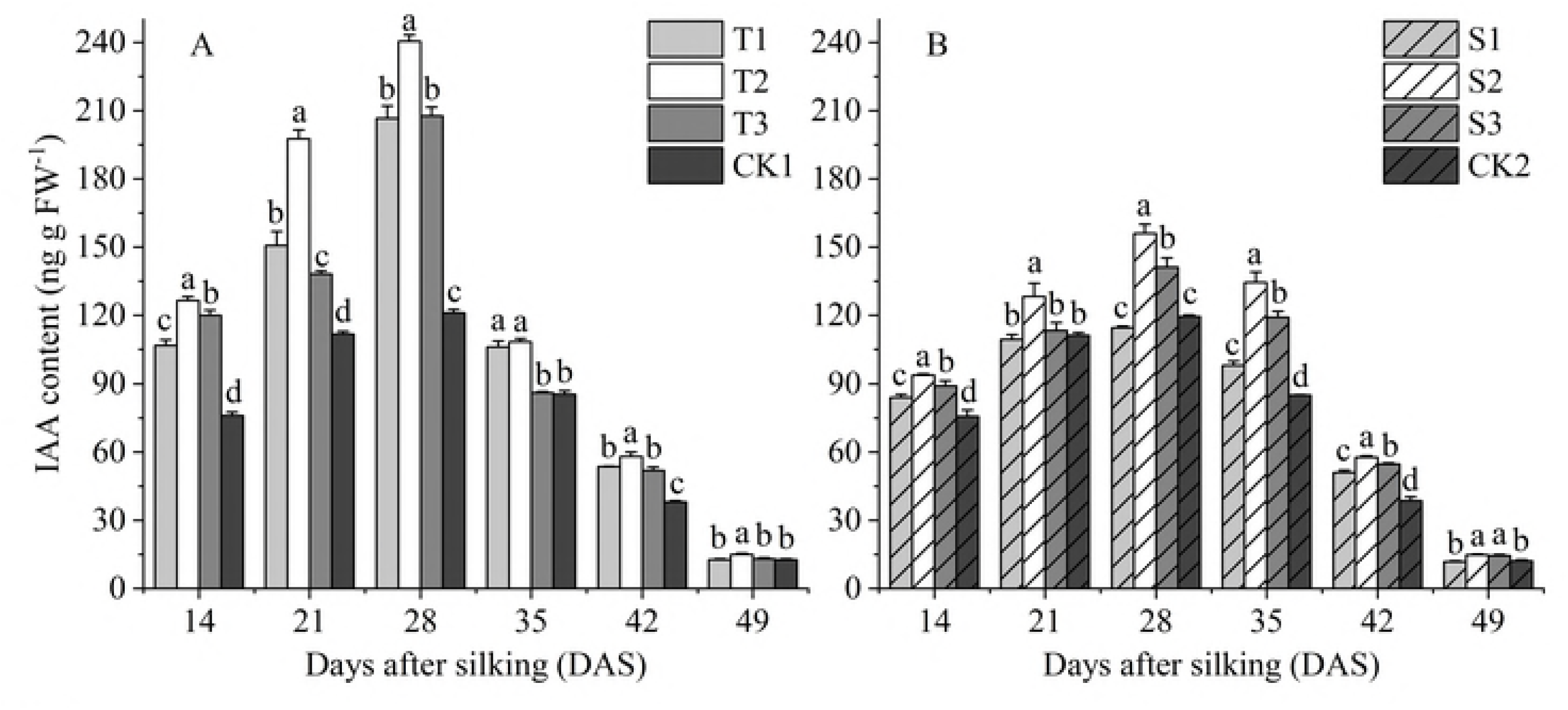
The effects of GA_4+7_ smearing application on IAA content in maize. CK1, T1, T2 and T3 represent shank-smearing with GA_4+7_ at the rate of 0, 10, 60, and 120 mg L^-1^, respectively. CK2, S1, S2 and S3 represent silk-smearing with GA_4+7_ at the rate of 0, 10, 60, 120 mg L^-1^, respectively. Different letters within each growth stage are significantly different (P < 0.05). The vertical bars represent the ± the standard error of the mean (n = 3).

**Fig. 4.**
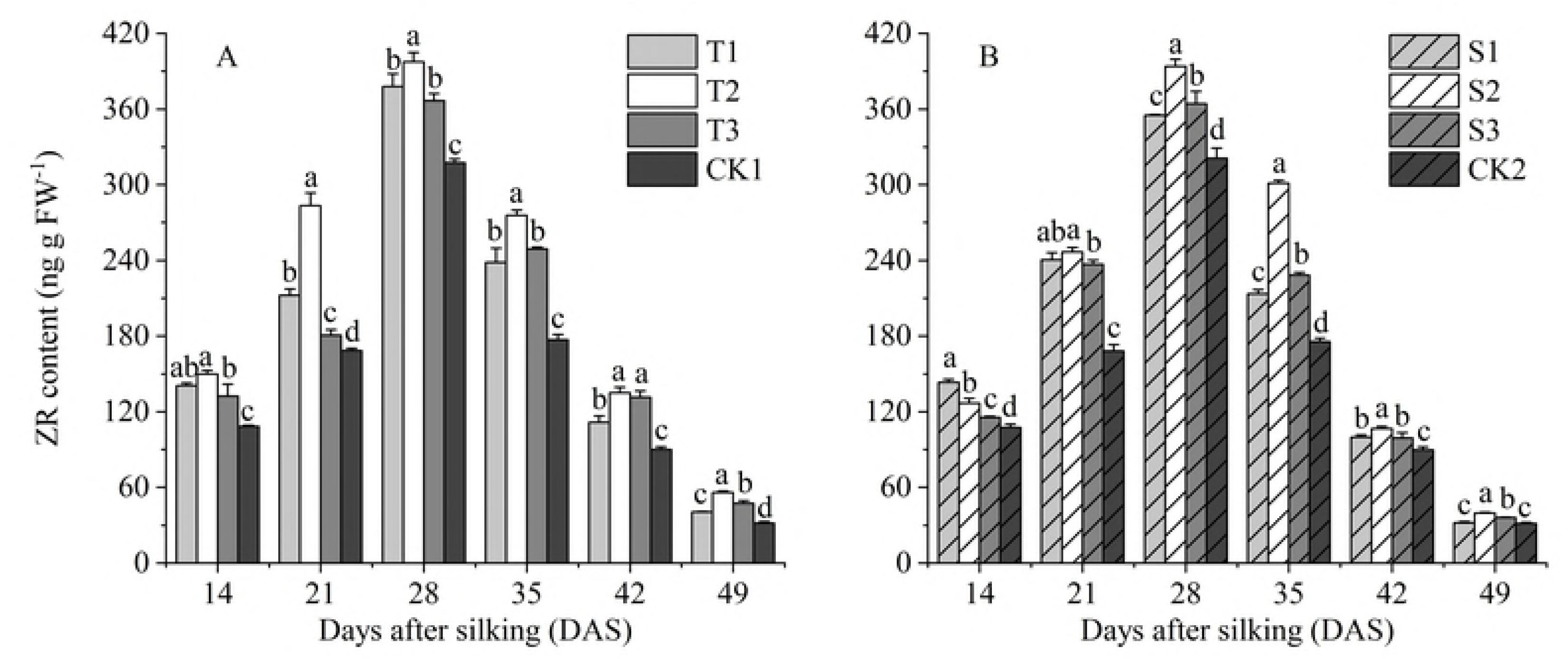
The effects of GA_4+7_ smearing application on ZR content in maize. CK1, T1, T2 and T3 represent shank-smearing with GA_4+7_ at the rate of 0, 10, 60, and 120 mg L^-1^, respectively. CK2, S1, S2 and S3 represent silk-smearing with GA_4+7_ at the rate of 0, 10, 60, 120 mg L^-1^, respectively. Different letters within each growth stage are significantly different (P < 0.05). The vertical bars represent the ± the standard error of the mean (n = 3).

#### 3.3.2 GA_3_ contents in grains

The GA_3_ contents in the grains exhibited a gradually decreasing trend during grain filling, except for S3 treatment (Fig. 5). S3 treatment decreased the level of GA_3_ in grains from 14 to 21 DAS and rose abruptly to a peak of 28 DAS, and then decreased steadily after the respective peak. The trend is different from other treatments, and it is necessary to deepen the research to explore the mechanism of its raise. Compared with control treatments, T2 and S2 treatments significantly increased the GA_3_ contents in grains and followed by T1 and S3. The average GA_3_ contents under T1, T2, and T3 treatments during the whole grain filling stage were increased by 9.1%, 26.4%, 5.1% compared with CK1, while that of S1, S2, and S3 treatments were greater by 11.6%, 25.9%, and 22.3% compared with CK2.

**Fig. 5.**
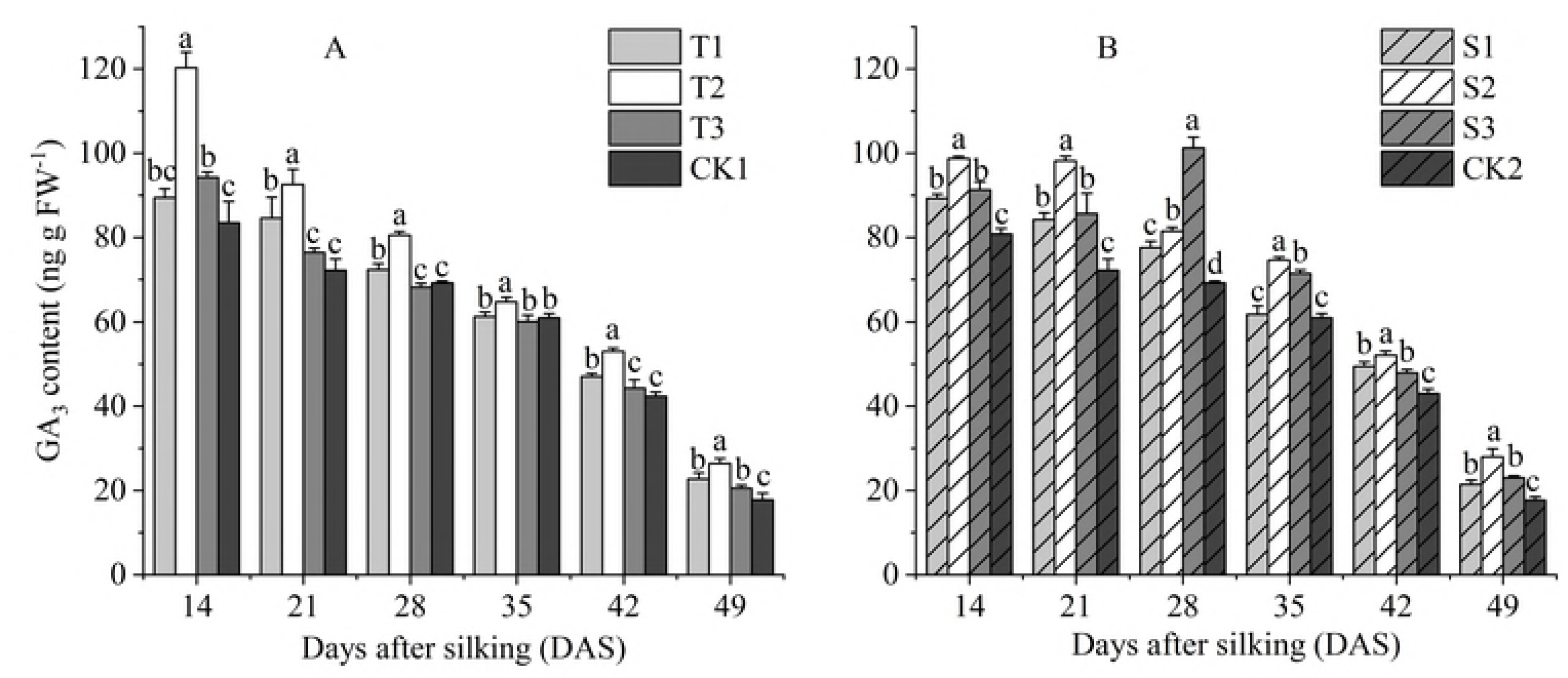
The effects of GA_4+7_ smearing application on GA_3_ contents in maize. CK1, T1, T2 and T3 represent shank-smearing with GA_4+7_ at the rate of 0, 10, 60, and 120 mg L^-1^, respectively. CK2, S1, S2 and S3 represent silk-smearing with GA_4+7_ at the rate of 0, 10, 60, 120 mg L^-1^, respectively. Different letters within each growth stage are significantly different (P < 0.05). The vertical bars represent the ± the standard error of the mean (n = 3).

#### 3.3.3 ABA contents in grains

The ABA content increased rapidly at the early grain-filling stage, reached to a maximum at 28 DAS, and then declined progressively in later stages. At the same sampling period, ABA content was decreased with the increase of GA_4+7_ concentration under shank-smearing treatments, while that under silking-smearing treatments was gradually increased with the increasing of GA_4+7_ concentration (Fig. 6). T1 and S3 treatments have the significant effects on ABA contents in grains at all sampling period, followed by S2 and T2. The average ABA contents during the whole grain filling stages under T1 and T2 treatments were increased by 14.0 ng g FW^-1^, 7.4 ng g FW^-1^, 12.8%, and 7.4% increases compared with CK1, whereas that of S2 and S3 increased by 7.9 ng g FW^-1^, 14.9 ng g FW^-1^, respectively, i.e., 13.6% and 7.9% increases compared with CK2. However, T3 and S1 had no significant effect relatively on increasing ABA contents compared with control treatment (CK).

**Fig. 6.**
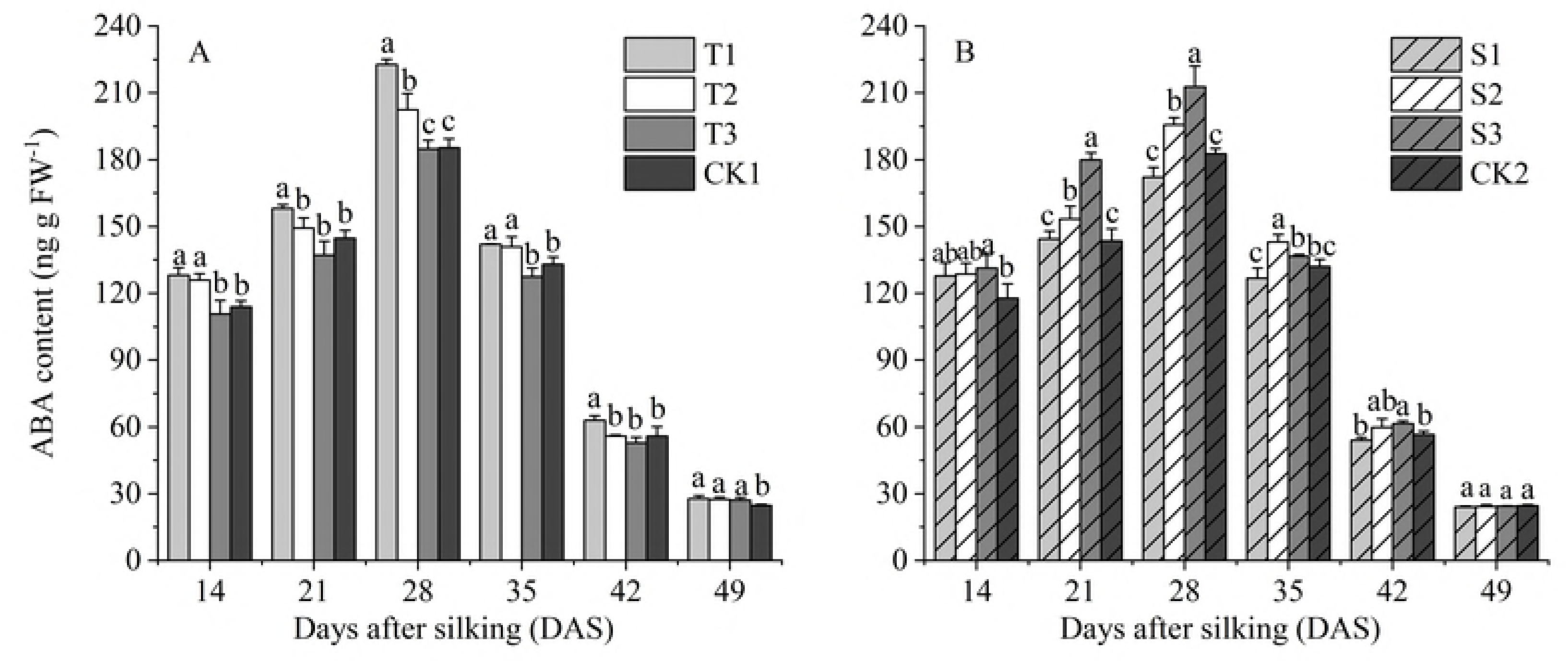
The effects of GA_4+7_ smearing application on ABA content in maize. CK1, T1, T2 and T3 represent shank-smearing with GA_4+7_ at the rate of 0, 10, 60, and 120 mg L^-1^, respectively. CK2, S1, S2 and S3 represent silk-smearing with GA_4+7_ at the rate of 0, 10, 60, 120 mg L^-1^, respectively. Different letters within each growth stage are significantly different (P < 0.05). The vertical bars represent the ± the standard error of the mean (n = 3).

### 3.4 Effects of GA_4+7_ on antioxidant enzymes

#### 3.4.1 SOD activity

The SOD activity showed a downward trend from 15 to 45 DAS under all treatments (Fig. 7). The SOD activity was significantly improved in GA_4+7_-treated plants at a varied level, Compared to control. 10 mg L^-1^ (T1 and S1) and 60 mg L^-1^ (T2 and S2) GA_4+7_ treatments had the significant effects for improving SOD activity than highest concentration of 120 mg L^-1^ GA_4+7_ (T3 and S3 treatments) and control. The SOD activity in T3 treatment was significantly greater than CK1 at 25 DAS, and no significant difference was associated with control at other stages. S3 treatment significantly increased the SOD activity compared with CK2 at 15 and 25 DAS; however, from 35 to 45 DAS, there was no significant difference between S3 and CK2. The SOD activity revealed an increase of 12.7%, 17.2% and 5.9% (mean of all sampling stages) in shank-smearing-treated plants, compared to CK1, while that of silk-smearing-treated plants was 12.3%, 16.6%, and 9.3% higher than CK2.

**Fig. 7.**
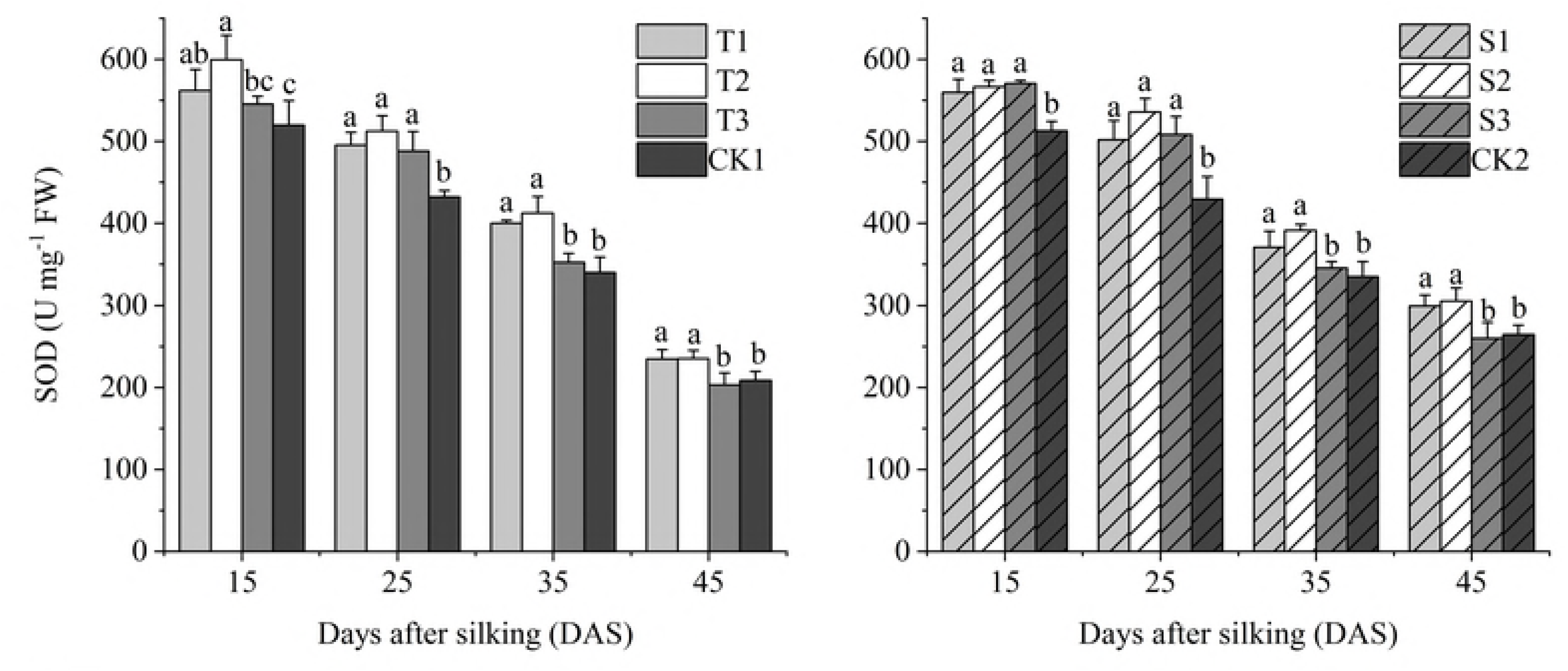
The effects of GA_4+7_ smearing application on SOD activity in maize. CK1, T1, T2 and T3 represent shank-smearing with GA_4+7_ at the rate of 0, 10, 60, and 120 mg L^-1^, respectively. CK2, S1, S2 and S3 represent silk-smearing with GA_4+7_ at the rate of 0, 10, 60, 120 mg L^-1^. Different letters within each growth stage are significantly different (P < 0.05). The vertical bars represent the ± the standard error of the mean (n = 3).

#### 3.4.2 POD activity

The POD activity in ear leaf displayed a gradually decreasing pattern of change from 15 to 45 DAS. And at the same stage, with the increase of GA_4+7_ concentrations, POD activity decreased gradually (Fig. 8). T1 treatment had the best effect on increasing the POD activity, followed by T2. There was no significant difference between T1 and T2. The POD activity revealed a significant increase of 9.0%, 45.6%, 10.8% and 15.8% in T1-treated plants, compared to CK1 at 15, 25, 35 and 45 DAS, while that of T2 was 6.1%, 37.4%, 14.2%, and 15% higher than CK1. Whereas at the early grain filling stage (from 15 to 25 DAS), the POD activity of T3 is higher than that of CK1, then the POD activity of T3 decreased rapidly, and at 45 DAS, the POD activity of T3 was significantly lower than that in CK1. The treatments of silk-smearing with GA_4+7_ significantly increased the POD activity at all sampling stages compared with CK2, except for T3 treatment at 15 DAS. At 15 and 25 DAS, S1 had the best effects on increasing the POD activity, followed by S2. And from 35 to 45 DAS, the POD activities of T1, T2, and T3 were not significantly different from each other. S1, S2, and S3 increased the average POD activities of the whole grain filling stages by 17.8%, 18.0% and 15.3%.

**Fig. 8.**
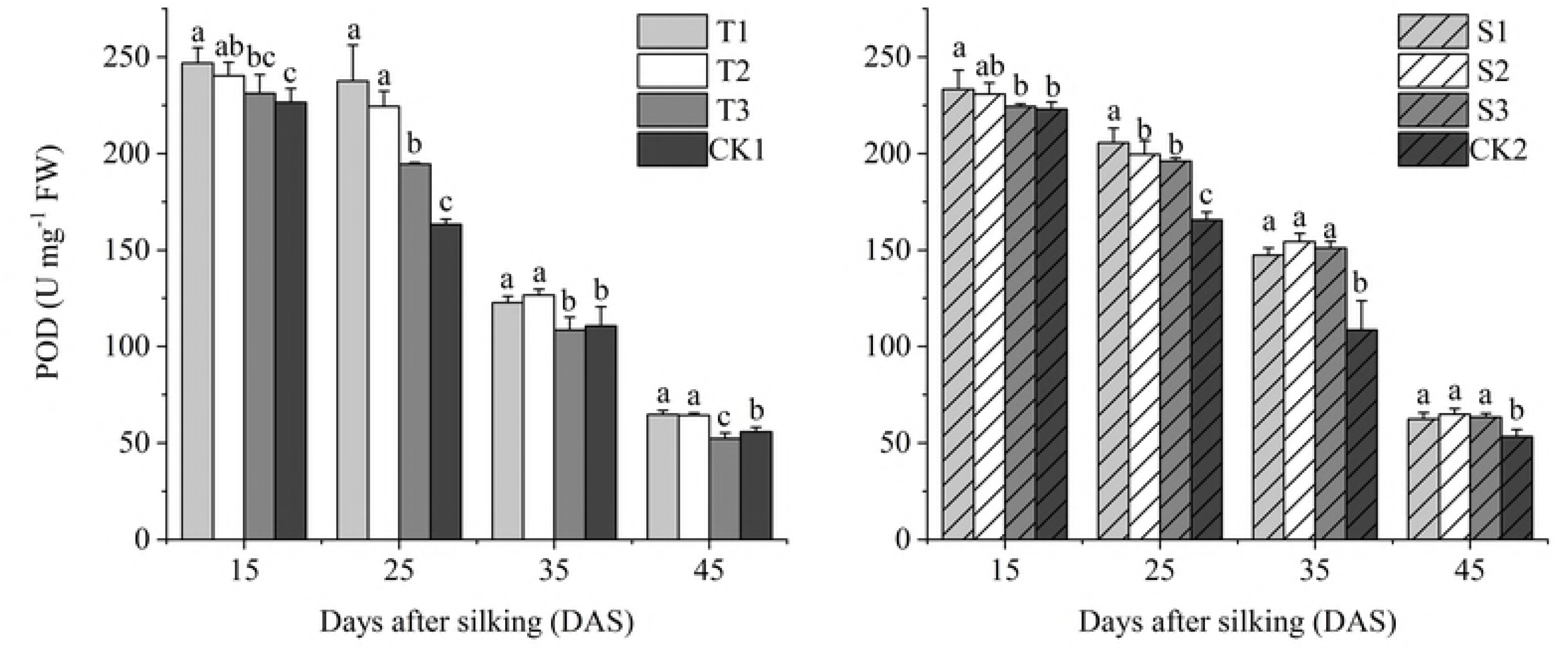
The effects of GA_4+7_ smearing application on POD activity in maize. CK1, T1, T2 and T3 represent shank-smearing with GA_4+7_ at the rate of 0, 10, 60, and 120 mg L^-1^, respectively. CK2, S1, S2 and S3 represent silk-smearing with GA_4+7_ at the rate of 0, 10, 60, 120 mg L^-1^. Different letters within each growth stage are significantly different (P < 0.05). The vertical bars represent the ± the standard error of the mean (n = 3).

#### 3.4.3 CAT activity

Application of GA_4+7_ had obvious effects on the activity of CAT and showed a gradually decreasing trend from 15 to 45 DAS (Fig. 9). With GA_4+7_ shank-smearing applications, T1 treatment showed the best effect of CAT activity at all sampling stages, followed by T2. And at 25 DAS, the CAT activity of T2 was significantly less than that of T1, but there was no significant difference in CAT activity between T1 and T2 in other sampling periods. The CAT activity in T3 was significantly higher than that of CK1 and significantly less than that of T2. All concentrations of GA_4+7_ silk-smearing treatments significantly increased CAT activity compared with CK2. The CAT activity of S2 was significantly higher than S1, S3, and CK2. The average of CAT activity of all sampling stages in T1, T2, and T3 treatments was increased by 58.4%, 46.1%, and 17.7% compared with CK1. Similarly, S1, S2, and S3 treatments improved the average CAT activity by 20.2%, 39.8%, and 24.8% compared with CK2.

**Fig. 9.**
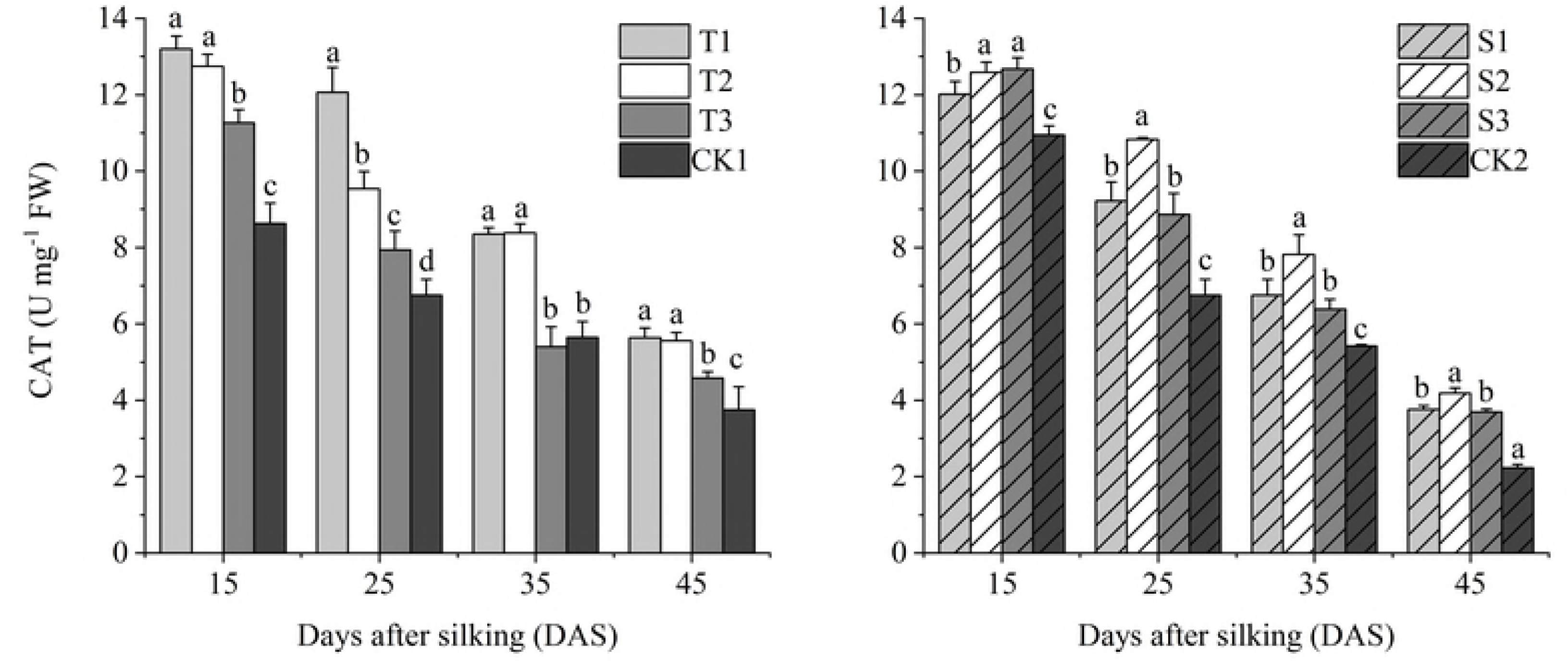
The effects of GA_4+7_ smearing application on CAT activity in maize. CK1, T1, T2 and T3 represent shank-smearing with GA_4+7_ at the rate of 0, 10, 60, and 120 mg L^-1^, respectively. CK2, S1, S2 and S3 represent silk-smearing with GA_4+7_ at the rate of 0, 10, 60, 120 mg L^-1^. Different letters within each growth stage are significantly different (P < 0.05). The vertical bars represent the ± the standard error of the mean (n = 3).

#### 3.4.4 MDA contents

The MDA content increased gradually from 15 to 45 DAS in all the treatments, with the increase in GA_4+7_ treatments significantly lower than that of the untreated control (Fig.10). All the silking-smearing treatments (S1, S2, and S3) had significant effects on decreasing the MDA from 15 to 45 DAS and were not significantly different from each other. While the T1 and T2 treatments significantly reduced the MDA contents at all sampling times and were not significantly different from each other. Moreover, T3 treatment exhibited relatively higher MDA contents than T1 and T2 treatments, and no significant difference was shown between T3 and CK1. The MDA content in T1 treatment was reduced by 11.8%, 11.7%, 19.0% and 11.5%, while that of T2 treatment was reduced by 11.9%, 10.0%, 19.5%, and 12.3% compared to CK1, at 15, 25, 35 and 45 DAS, respectively. And the average MDA content from 15 to 45 DAS in S1, S2, and S3 treatment were reduced by 1.36 U mg^-1^ FW, 1.43 U mg^-1^ FW, 1.22 U mg^-1^ FW, 11.5%, 12.0% and 10.2% reduced compared with CK2.

**Fig. 10.**
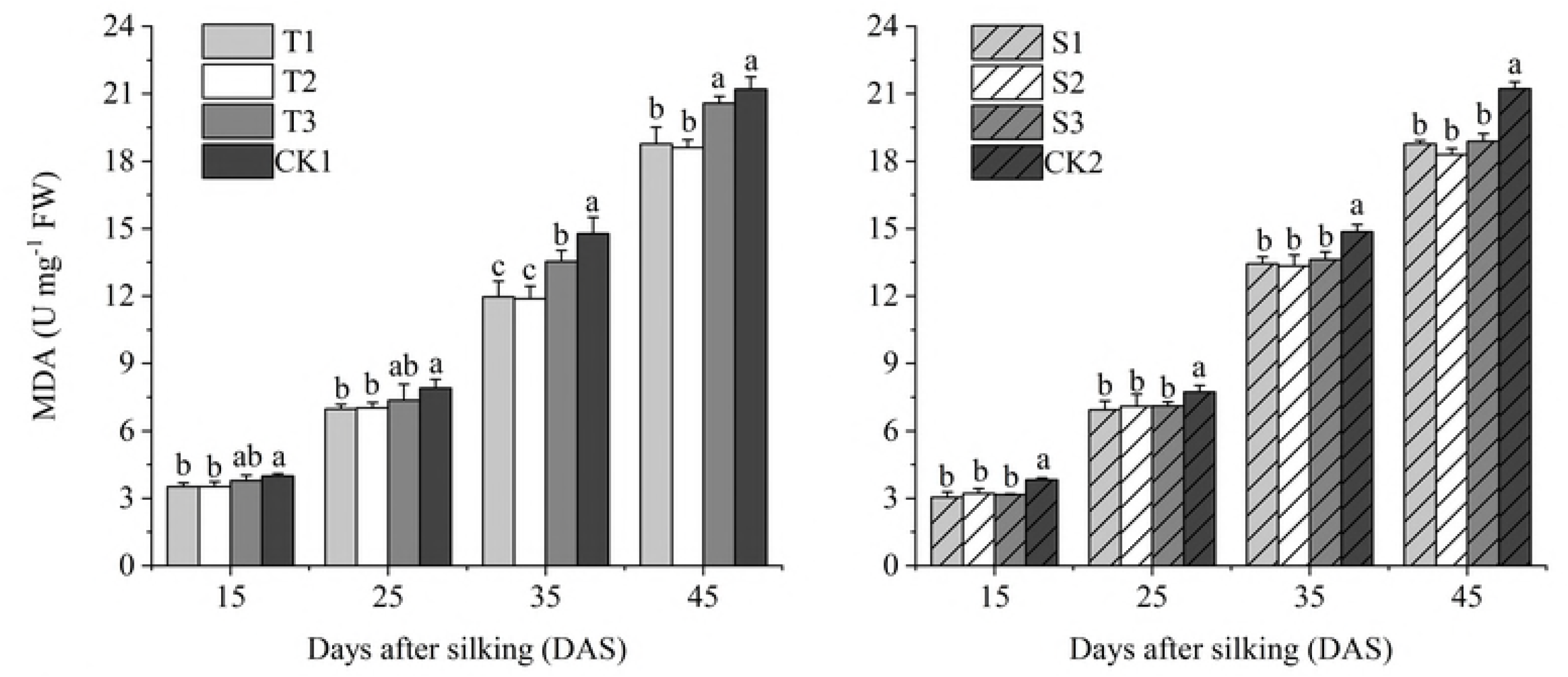
The effects of GA_4+7_ smearing applications on MDA content in maize. CK1, T1, T2 and T3 represent shank-smearing with GA_4+7_ at the rate of 0, 10, 60, and 120 mg L^-1^, respectively. CK2, S1, S2 and S3 represent silk-smearing with GA_4+7_ at the rate of 0, 10, 60, 120 mg L^-1^, respectively. Different letters within each growth stage are significantly different (P < 0.05). The vertical bars represent the ± the standard error of the mean (n = 3).

## 4 Discussion

### 4.1 Effects of GA_4+7_ smearing application on grain yield

The present study illustrates the positive effects of GA_4+7_ on grain yield of high-density maize by regulating the endogenous hormones level in grains and the activities of antioxidant enzymes in leaves to promote the grain filling rate and regulate the leaves senescence. Previously, GA_4+7_ have been shown to be an effective growth regulator for several crops that could increase the yield and improve the income for farmers [33-35]. In our study, GA_4+7_ at a rate of 10 (T1) and 60 (T2) mg L^-1^ of under shank-smearing application significantly increased the grain yield of maize in 2015 and 2016. With the silk-smearing application, all concentrations of GA_4+7_ significantly improved the maize yield in both 2015 and 2016. The application of GA_4+7_ improved the agronomic characteristics of ears, increased the kernel number per ear and thousand kernels weight. The yield potential of maize can be divided into three key components: kernel number per ear, grain weight, and a number of ear per plant. However, the average thousand grain weight under shank-smearing treatments in 2015 and 2016 were higher than that in silk-smearing treatments, and the average kernel number of silk-smearing treatments in 2015 and 2016 were higher than that in shank-smearing treatments. These results indicated that GA_4+7_ with shank-smearing application improved the yield mainly by increased the grain weight, while silk-smearing with GA_4+7_ increased the kernel number per ear to promote the maize yield.

### 4.2 Relationship of hormone changes and maize grain filling

Cytokinins (CTKs) plays a significant role in the regulation of grain filling process, and high CTK levels in maize spike tissues are markedly associated with kernel development [44-45]. CTKs are generally found in the endosperm of developing seeds and may be required for cell division during the early phase of seed setting in other cereal crops [44, 46-49]. Besides CTKs, IAA also plays a significant role in regulating grain filling [50]. Xu, et al. [24] and Fu, et al. [51] suggested that regulation of the IAA content in grains could potentially increase the weight of inferior grains. In present study, ZR and IAA contents in the maize grains showed a significantly positive correlation with the maximum grain weight, the maximum and mean grain-filling rates (Table 4). The results indicated that ZR and IAA contents in grains were involved in regulating maize grain filling. In addition, the changes of ZR and IAA contents in grains during grain filling stages showed a similar pattern. The ZR and IAA contents in the grains transiently increased in the early grain-filling stage and then decreased, reaching a maximum at 28 DAS. We also observed that the IAA and ZR contents between GA_4+7_-treated plants and untreated maize had significant difference at early filling stage. However, with the development of grain filling process, the difference of IAA and ZR contents between GA_4+7_-treated maize and control gradually narrowed (Fig. 3 and 4). The researchers suggested that auxin and CTKs control endosperm cell growth and division [24, 52]. Seth [53] and Singh [54] stated that high IAA levels in a sink organ can create an "attractive power", leading to increased cytokinins levels in grains. These previous findings as well as the findings in this research indicate that IAA and ZR contents may control early grain filling in maize.

**Table 4.**
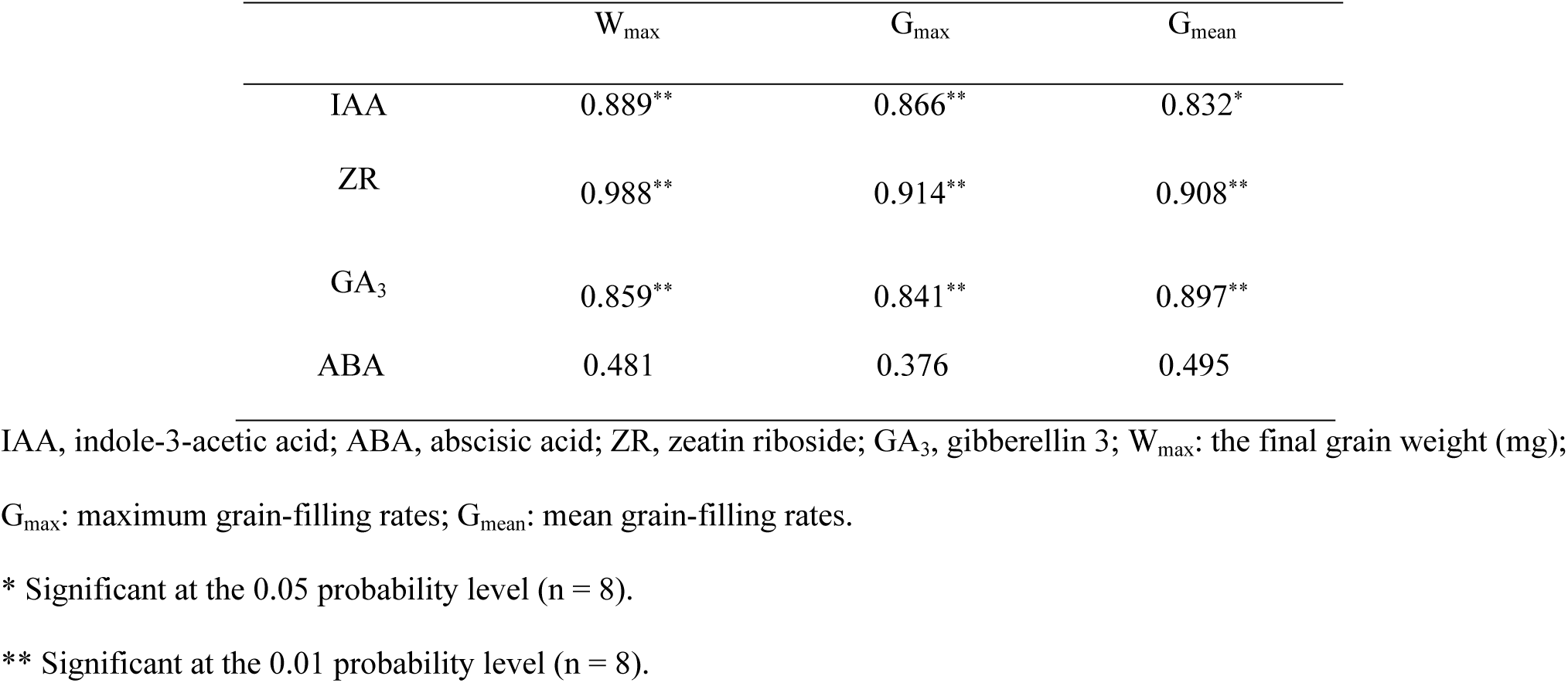
Correlation coefficients of mean hormone contents in maize grain with the maximum grain filling rate (G_max_), mean grain filling rate (G_mean_), and maximum grain weight (W_max_) of maize

Furthermore, ABA and GAs also play important roles in regulated grain filling. Yang, et al. [27] suggested that a higher ABA concentration in grains enhanced the remobilization of prestored carbon to the grains and accelerated the grain filling rate. And the previous studies showed that ABA content in grains was significantly and positively correlated with the maximum and mean grain filling rates and maximum grain weight [25-26, 55]. In our present study, the ABA content in grains were increased by exogenous GA_4+7_, and there was positively correlated between ABA and grain filling rate (Table.4). However, the correlation between GAs and grain filling rate and grain weight were not in consistent. Liu, et al. [25] and Liu, et al. [55] suggested that the content of GAs in the grains was not significantly correlated with the maximum grain weight or the maximum and mean grain-filling rates. In contrast, Our findings are consistent with Ali, et al. [26], that GAs was significant correlated with the maximum grain weight and the maximum and mean grain-filling rates. Yang, et al. [56] stated that spraying exogenous GA reduced the ratio of ABA to GA in the grains. These results indicated that the application of exogenous GA_4+7_ changes the content of ABA and GA in the grains, and a hormonal balance, rather than individual hormone content, regulates and improves the maize grain filling.

### 4.3 Effects of GA_4+7_ smearing application on the antioxidant enzymes

Reactive oxygen species (ROS) are continually being produced in plants, which can act as a signal molecule in plants and trigger a series of cellular responses [36]. In plants, an increase in the ROS levels exceeding the detoxification levels of plant tissues could be toxic. However, plants have evolved an enzymatic and non-enzymatic antioxidants defense mechanism to effectively scavenge the ROS and maintain a proper balance within the plants [57]. Enzymatic antioxidants mainly include superoxide dismutase (SOD) peroxidase (POD) ascorbate peroxidase (APX), and catalase (CAT). However, senescence and various environmental stresses could disrupt the balance between ROS generation and detoxification, resulting in lipid peroxidation, chlorophyll degradation and loss of cell membrane integrity [58]. Indeed, our result showed a remarkable increase in the activities of antioxidant enzymes including SOD, POD, and CAT, and decreased MDA accumulation in the GA_4+7_-treated maize plants at various growth stages. These trends could be interpreted as enhanced scavenging capacity of reactive oxygen species and reduced membrane lipid oxidation of treated plants. It is well-known that chloroplasts are a potential source for generation of active oxygen [59]. Therefore, GA_4+7_-treated plants kept more photosynthetic pigments content and subsequent in the delayed aging.

With the knowledge of that developing seeds contain large amounts of hormones, which possess the ability of inducing directional movement of nutrients within plants. The leaves senesce appears when a "sink" (young developing leaves or seeds) needs the nutrients which were moved from the rest of the plant. Seth and Wareing [60] suggested that hormone-directed transport plays an important role in directing the movement of nutrients towards developing seeds. These indicated that the application of exogenous GA_4+7_ changed the level of hormones in seeds, thus affecting the antioxidant enzymes in leaves and the senescence of maize. However, there are little research about exogenous GA_4+7_ on reactive oxygen metabolism in maize, and the specific mechanism needs further study.

## 5 Conclusions

Application of GA_4+7_ under high density leads to an increase in the level of IAA, ZR, GA_3,_ and ABA. This increase is attributed to improving the grain filling rate, grain weight and the number of kernels ear^-1^ were also increased, as well as increased the yield. In addition, GA_4+7_ applications improved the activities of antioxidant enzymes in leaves senescence. The grain filling period could be prolonged by delaying the senescence of maize so that the grain filling was sufficient and eventually increased the yield. Our results showed the positive effect of GA_4+7_ on maize grain filling rate and antioxidant enzymes, and could effectively be used for crop improvement, especially for cereal crops. At the rate of 60 mg L^-1^, GA_4+7_ showed the greatest effect for shank-smearing and silk-smearing (T2 and S2), followed by 10 mg L^-1^ (T1) for shank-smearing and 120 mg L^-1^ (S3) for silk-smearing. The results from the present study illustrate that GA_4+7_ application could efficiently be used for altering the level of hormones in grains and antioxidant enzymes in ear leaf, would be useful for enhancing grain filling rate and delaying leaves senescence and increasing grain yield of maize.

